# Implementation of Genomic Variant Calling: A Novel Approach

**DOI:** 10.1101/2020.05.31.126144

**Authors:** Ambarish Kumar, Ali Haider Bangash

## Abstract

Genomics has emerged as one of the major sources of big data. The task of augmenting data-driven challenges into bioinformatics can be met using technologies of parallel and distributed computing. GATK4 tools for genomic variants detection are enabled for high-performance computing platforms – SPARK Map Reduce framework. GATK4+WDL+CROMWELL+SPARK+DOCKER is proposed as the way forward in achieving automation, reproducibility, reusability, customization, portability and scalability. SPARK-based tools perform equally well in genomic variants detection with that of standard implementation of GATK4 tools over a command-line interface. Implementation of workflows over cloud-based high-performance computing platforms will enhance usability and will be a way forward in community research and infrastructure development for genomic variant discovery.

## 1 Introduction

Much required and immediate felt need of the genomics research, especially variant detection, is a customised pipeline for detection of genomic variants. There are various tools for genomic variant detection whose performances vary in comparison to one another. GATK4[2] tools use convolutional neural networks – CNN for genomic variants detection. Along with GATK4, WDL – Workflow Description Language [2], CROMWELL [2] and DOCKER caters the need for building customised pipelines or workflows for genomic variant discovery. Moreover GATK4 tools are enabled for SPARK MapReduce framework to achieve computational scalability. Workflows hosted over high-performance cloud-based computing infrastructure will meet the emerging trend and technology towards virtual laboratory and infrastructure enhancement for genomic variant discovery.

## 2 Materials & Methods

Entire setup is run over the Linux operating system and JAVA 1.8 environment. Running GATK4 in cluster mode requires setting SPARK cluster and HADOOP installation. There is no version-specific requirement or incompatibility among software except specific JAVA 1.8. All of the required software and their installation for performing the study are tabulated in Table-01 and Table-02 of supplementary material. Steps for multi-node cluster setup are mentioned in the SPARK Cluster Setup section of supplementary material.

### 2.1 Standard operating protocol to simulate RNASEQ reads using RSEM

RSEM [1] contains a simulator to simulate RNASEQ reads. It generates single-end or paired-end RNASEQ reads in fasta or fastq format. Protocol is designed using bowtie as an aligner. In addition, samtools is required as a dependency. Example of command-line implementation is shown for simulating paired-end ebola RNASEQ reads in fastq format.

### 2.2 Setting path environment

export PATH=$PATH:/absolute path to bowtie executables/

### 2.3 Preparing Reference Sequences

Prepare reference transcripts for RSEM and optionally build bowtie indices of reference genome.

#### Command-line usage

rsem-prepare-reference <reference_fasta_file> --bowtie2 <reference_name>

#### Command-line arguments

- rsem-prepare-reference = RSEM command to prepare reference transcripts
- <reference_fasta_file> = reference transcript fasta file (from which one has to simulate RNASEQ reads).
- --bowtie2 = option to build bowtie indices.
- <reference_name> – base name for the reference.

#### Example

$ rsem-prepare-reference ebola.fasta --bowtie2 ebola

### 2.4 Calculating Expression Values

Following calculates the expression level of transcripts using real sequenced RNASEQ reads:

> rsem-calculate-expression

It also learns all of the sequencing parameters and builds a model file. That model file is then used by rsem-simulator to assign sequencing parameters to simulated reads.

#### Command-line usage

rsem-calculate-expression --paired-end <upstream_read_file> <downstream_read_file> <reference_name> <sample_name>

#### Command-line arguments

- rsem-calculate-expression = RSEM command to estimate gene and isoform expression from RNA-Seq data level and generate model file based on the expression pattern
- >upstream_read_file> = real sequenced RNASEQ reads generated from forward strand i.e left.fq reads
- <downstream_read_file> = real sequenced RNASEQ reads generated from reverse strand i.e right.fq reads
- <reference_name> = The name of the reference used
- <sample_name> = The name of the sample analyzed (Note that all of the output files are prefixed by this name)

#### Example

$rsem-calculate-expression --paired-end ebola_ref_reads1.fq ebola_ref_reads2.fq ebola ebolaRNA

### 2.5 Simulation

RSEM contains a simulator to simulate RNA-Seq data based on parameters learned from real data sets.

#### Command-line usage

$rsem-simulate-reads <reference_name> <estimated_model_file> <estimated_isoform_results> <theta0> N <output_name>

#### Command-line arguments

- <reference_name> = The name of RSEM references
- <estimated_model_file> = This file describes how the RNA-Seq reads will be sequenced given the expression levels
- <estimated_isoform_results> = This file contains expression levels for all of the isoforms recorded in the reference.
- <theta0> = Option for fraction of reads sequenced from noise.
- N = Option for total number of reads to be simulated.
- <output_name> – Prefix for all output files.

#### Example

$rsem-simulate-reads ebola ebolaRNA.stat/ebolaRNA.model ebolaRNA.isoforms.results 0.2 10000 ebolareads

The final output consists of the fastq format paired-end reads appended with _1 and _2 in order to denote the forward and reverse paired ends.

- Left-end reads = ebolareads_1.fq
- Right-end reads = ebolareads_2.fq

### 2.6 Ebola test dataset generation

#### Test dataset containing non-structural variants

To benchmark the performance of each workflow, paired-end RNASEQ reads are simulated from manually mutated Ebola reference genome with 10 SNPs, 10 small INDELs, 05 INVERSIONs and their reverse complement as well as 05 TRANSLOCATIONs [dataset][dataset1].

##### SNPs

Mutated genomic bases to call SNPs and their detection status by GATK4 are tabulated in Table-03 of supplementary material.

##### INDELs

Mutated genomic bases to call INDELs and their detection status by GATK4 are tabulated in Table-04 of supplementary material.

##### TRANSLOCATIONs

Translocated genomic bases are tabulated in Table-05 of supplementary material.

##### INVERSIONs

Called inversions and their reverse complement in the mutated Ebola genome are tabulated in Table-06 of supplementary material.

Description is tabulated into Table-07 of supplementary material.

#### Test dataset containing structural variants

Similarly, the reference ebola genome is manually mutated to call structural variants. Structural variant calling is performed over two datasets. The first dataset contains seven pairs of simulated RNASEQ fastq reads, each one simulated from separate mutant ebola genome containing one large deletion, one insertion, one duplication, one translocation, one inversion, one complex variant1 (INSDUPDEL) and one complex variants2 (DELDUPDEL) respectively. The second dataset comprises of seven pairs of simulated RNASEQ fastq reads, each one simulated from separate mutant ebola genome containing two large deletions, two insertions, two duplications, two translocations, two inversions, one complex variant1 (INSDUPDEL) and one complex variants2 (DELDUPDEL) respectively [dataset][dataset2][dataset3].

The description of mutated genomic bases in both of the datasets is tabulated into Table-08 and Table-10 of supplementary material. The lists of both the simulated ebola datasets used for structural variant calling are provided in Table-09 and Table-11 of supplementary material.

### 2.7 Writing WDL script

WDL script is block-structured and consists of following components or blocks – workflow, task, call, command, output. Docker-based workflow has an additional block of runtime within the task block specifying runtime environment condition. Defining the workflow is the first most step in writing the WDL script. Within the workflow block, workflow level input variables are defined and there exists a call block for task call and for the enumeration of workflow level as well as task level input variables. Task call block contains all input variables necessary for running the task. It is defined outside the workflow block and it contains defined task level input variables, command block, output block and runtime block within it. [WDL scripts].

#### Adding input variables

There are two levels of input variables:

– task level
– workflow level

Both types of variables follow the same rule of declaration. These variables serve as placeholders and aliases for the actual filename and parameter values.

#### Adding workflow level variables

Workflow level input variables are available to any of the tasks that it calls.

#### Adding task level variables

These input variables are declared at the top of the task block. Code block within the call function contains input variables required for running the task as well as enumeration of workflow level variables and task-level variables.

Command block resides within the task block and specifies the literal command line to run with placeholders for the variable parts of the command line that need to be filled in.

#### Plumbing of tasks

Also referred to as *‘linear chaining of tasks’* as it is used to form a sophisticated pipeline or workflow, plumbing is done by connecting tasks through their inputs and outputs. WDL allows the output of any task to be referred to within the call statement of another task.

Linear chaining of tasks into SNPs and INDELs detection workflows are as follows:

> Aligment→AddOrReplaceReadGroups→SortSam→ReferenceSeqIndex→ ReferenceSeqDictionary→MarkDuplicates→HaplotypeCaller→ VariantFilteration→SelectSNPs→SelectINDELs

Linear chaining of tasks into structural variants detection workflows are as follows:

> Alignment→AddOrReplaceReadGroups→SortSam→ReferenceSeqIndex→ReferenceSeqDictionary→BwaMemIndexImageCreator→FindBadGenomicKmers→FindBreakpointEvidence→ SvDiscovery

Sequential execution of tasks depends upon the DAG graph of tasks generated based on passed-on input variables to task command. One need not worry about the order or sequence of task call or task definition into his or her WDL script.

Detailed descriptions for writing WDL scripts are discussed in the Writing WDL Script section of supplementary material.

There are six WDL scripts [WDL scripts] proposed into the study corresponding to each workflow for genomic variants discovery.

Table-12.0 of supplementary material lists all six workflows with their WDL script and JSON format input file.

### 2.8 Validate WDL script syntax

The ‘validate’ function of WDL tool parses the WDL script for the syntax correction.

#### Command-line usage

$ java −jar womtool.jar validate <workflow.wdl>

#### Command-line arguments

- womtool.jar = womtool java jar file
- validate = option to validate WDL script
- <workflow.wdl> = WDL script corresponding to workflow

### 2.9 Specify WDL script inputs

The input function of the WDL tool generates a template of inputs for the WDL script into JSON format.

#### Command-line usage

$ java −jar womtool.jar inputs <workflow.wdl> > <workflow_inputs.json>

#### Command-line arguments

- womtool.jar = womtool java jar file
- Inputs = option to generate inputs
- <workflow.wdl> = WDL script corresponding to workflow
- <workflow_inputs.json> = generated JSON format input file

workflow_inputs.json lists all of the inputs to all the tasks present into the WDL script. Inputs are present into the key-value pair.

### 2.10 CROMWELL Execution

Cromwell executes in two modes:

- run mode
- server mode

#### ‘Run’ mode

In the run mode, WDL script executes over command-line interface and exits when the workflow completes.

##### Command-line usage for running WDL script in the run mode

$ java −jar cromwell.jar run <workflow.wdl> --inputs <workflow_inputs.json>

##### Command-line arguments

- cromwell.jar = Java jar file for Cromwell
- run = option for the run mode
- <workflow.wdl> = WDL script
- --inputs = option for input
- <workflow_inputs.json> = JSON format input file

#### ‘Server’ mode

In the server mode, WDL script executes over REST API.

##### Command-line usage for running WDL script over REST API

$ java −jar cromwell.jar server

##### Command-line arguments

- cromwell.jar = Java jar file for Cromwell
- server = option to start Cromwell REST API

The above command starts the Cromwell web server. Following are additional functionalities:

- “POST” option is to submit a workflow for execution.
- “Try it out” parameter enables options for browsing the workflow – WDL script and input file – from the local machine.
- “Execute” option is to execute the workflow.

The submission status and workflow_id is contained within the Response body over the REST API.

Workflow timing diagram can be observed at http://localhost:8000/api/workflows/v1/workflow_id/timing.

Each workflow uses bowtie as an aligner. PATH environment variable to access bowtie executables is set into the “Alignment” task of each workflow.

It should be noted that prior to running the workflows which are not using DOCKER as run-time, one has to build gatk local jar after running the following command:

$ export GATK_LOCAL_JAR=<absolute_path_to_local_gatk_jar_package>

*<absolute_path_to_local_gatk_jar_package> is the absolute path of the gatk jar file placed on the local machine.*

## 3 Results

### 3.1 Analysis of variant calling method performance

#### SNPs detection

**Table-01.** SNPs detection performance analysis

| Predicted condition |  |  |  |  |
| --- | --- | --- | --- | --- |
| Condition |  | Positive | Negative |  |
|  | Positive | True positive<br>08 | False negative<br>02 | Sensitivity<br>(TPR)<br>0.8 |
|  | Negative | False Positive<br>03 | True negative | FNR<br>0.2 |
|  |  | Precision<br>0.8 | FDR<br>0.2 |  |

#### INDELs detection

**Table-02.** INDELs detection performance analysis

INDELs detection **Table-02.** INDELs detection performance analysis
| Predicted condition |  |  |  |  |
| --- | --- | --- | --- | --- |
| Condition |  | Positive | Negative |  |
|  | Positive | True positive<br>04 | False negative<br>06 | Sensitivity<br>(TPR)<br>0.4 |
|  | Negative | False Positive<br>03 | True negative | FNR<br>0.6 |
|  |  | Precision<br>0.6 | FDR<br>0.4 |  |

#### SVs

Structural variants detection across both the datasets are tabulated and summarized as follows:

**Table-03.** Structural variants detection status across dataset-1

| Mutation type |  | Detection Status |
| --- | --- | --- |
|  | Deletion | Detected |
|  | Insertion | Not detected |
|  | Inversion | Not detected |
|  | Duplication | Not detected |
|  | Translocation | Detected |
| CPX1 - INSDUPDEL | Insertion | Detected |
|  | Duplication | Not detected |
|  | Deletion | Detected |
| CPX2 - DELDUPDEL | Deletion | Detected |
|  | Duplication | Not detected |
|  | Deletion | Detected |

**Table-04.** Structural variants detection status across dataset-2

| Mutation type |  | Detection status |
| --- | --- | --- |
|  | Deletion1 | Detected |
|  | Deletion2 | Detected |
|  | Insertion1 | Not detected |
|  | Insertion2 | Not detected |
|  | Inversion1 | Not detected |
|  | Inversion2 | Not detected |
|  | Duplication1 | Not detected |
|  | Duplication2 | Not detected |
|  | Translocation1 | Detected |
|  | Translocation2 | Detected |
| CPX1 - INSDUPDEL | Insertion | Not detected |
|  | Duplication | Not detected |
|  | Deletion | Detected |
| CPX2 - DELDUPDEL | Deletion | Detected |
|  | Duplication | Not detected |
|  | Deletion | Detected |

Table-03 and Table-04 summarize GATK4 structural variants detection performance. One can observe that:

- All large DELETIONs are detected with 100% accuracy and precision.
- All TRANSLOCATIONs are detected with 01 FP into dataset-1 and 02 FP into dataset-2.
- INVERSIONs and DUPLICATIONs remain completely undetected across both of the datasets.
- Only one large novel sequence INSERTION is detected i.e of CPX1 – INSDUPDEL of dataset1.

Implementation of GATK4 tools over SPARK MapReduce framework has similar performance – in terms of genomic variants detection – like that of standard implementation over a non-SPARK platform.

Entire run for the execution of workflows containing SPARK based tools is set over a multi-node SPARK cluster configured with single-core virtual-machines – master node and slave node. To achieve enhanced scalability into computational time one may increase the number of allocated CPUs or cores to the SPARK based tools. -- spark-master local [] option as a command-line argument allows us to allocate an increased number of CPUs or cores to the running SPARK based GATK4 tools.

## 4 Discussion

With improved functionalities and features, GATK4 uses leading-edge machine learning techniques – CNN to detect genomic variants. Written WDL scripts for genomic variants detection workflows provide automation, reproducibility, reusability, customization, portability, scalability and hence extensive performance gain to users, especially biological researchers working into the area. It bridges the skills gap between non-expert users and high-performance computational platforms. These workflows can be uploaded to a cloud-based high-performance computing platform as virtual infrastructure for community research. Nonetheless, this study will help in accelerating the pace of genomics research and a way forward towards a built-in solution and enhanced infrastructure for biological researchers working in the area.

## References

[WDL scripts] Ambarish Kumar. (2019, September 13). Implementation of Genomic Variant Calling Using GATK4, SPARK, WDL, CROMWELL and DOCKER Over Simulated Ebola NGS Dataset. Zenodo. http://doi.org/10.5281/zenodo.3407838

[dataset] Jawaharlal Nehru University, New Delhi, India (2019). Implementation of Genomic Variant Calling Using GATK4, SPARK, WDL, CROMWELL and DOCKER Over Simulated Ebola NGS Dataset. Raw sequence reads. SRA submission. NCBI BioProject Accession number : PRJNA564447 and BioProject ID: 564447

[dataset1] Ambarish Kumar. (2020). Implementation of Genomic Variant Calling Using GATK4, SPARK, WDL, CROMWELL and DOCKER Over Simulated Ebola NGS Dataset. [Data set]. Zenodo. http://doi.org/10.5281/zenodo.3840885

[dataset2] Ambarish Kumar. (2020). Implementation of Genomic Variant Calling Using GATK4, SPARK, WDL, CROMWELL and DOCKER Over Simulated Ebola NGS Dataset. [Data set]. Zenodo. http://doi.org/10.5281/zenodo.3840893

[dataset3] Ambarish Kumar. (2020). Implementation of Genomic Variant Calling Using GATK4, SPARK, WDL, CROMWELL and DOCKER Over Simulated Ebola NGS Dataset. [Data set]. Zenodo. http://doi.org/10.5281/zenodo.3840897

[1] Li B, Dewey CN. RSEM: accurate transcript quantification from RNA-Seq data with or without a reference genome. BMC Bioinformatics. 2011;12:323. Published 2011 Aug 4. doi:10.1186/1471-2105-12-323

[2] Voss K, Gentry J and Van der Auwera G. Full-stack genomics pipelining with GATK4 + WDL + Cromwell [version 1; not peer reviewed]. F1000Research 2017, 6(ISCB Comm J):1379 (poster) (doi: 10.7490/f1000research.1114631.1).

[3] Zaharia, Matei, et al. “Spark: Cluster computing with working sets.” HotCloud 10.10-10 (2010): 95.

