## Supplementary material for "Implementation of Genomic Variant Calling: A Novel Approach"

Ambarish Kumar ^1[0000-0002-4923-046X]^ and Ali Haider Bangash^2[0000-0002-8256-3194]^

^1^ School of Computational and Integrative Sciences, Jawaharlal Nehru University, New Delhi, India

^2^ Shifa College of Medicine, STMU, Pakistan


**Supplementary Materials**

1 **Materials and Methods**
  1.1 **Software Setup Download & Installation**

Software setup download and installation are tabulated below.

**Table-01.** Software setup download

| Software | Version | Setup download |
| --- | --- | --- |
| JAVA | 8 | [www.oracle.com/technetwork/java/javase/](http://www.oracle.com/technetwork/java/javase/)  downloads/index.html |
| WOMltool | 50 | https://github.com/broadinstitute/cromwell/ |
| CROMWELL | 50 |  |
| GATK4 | 4.1.7.0 | https://github.com/broadinstitute/gatk/releases |
| SPARK | 2.4.3 | spark.apache.org/downloads.html |
| HADOOP | 2.6.5 | https://hadoop.apache.org/releases.html |
| GATK4 Docker container image | 4.1.7.0 | https://hub.docker.com/r/broadinstitute/gatk/ |
| Bowtie | 2.4.1 | https://sourceforge.net/projects/bowtie-bio/files/bowtie2/2.4.1 |
| Virtual-Box | 6.0 | www.virtualbox.org |
| Ubuntu OS | 16.04 LTS | ubuntu.com/download |
| R | 3.6.1 | cran.r-project.org |
| RSEM | 1.3.1 | deweylab.github.io/RSEM/ |
| Samtools | 1.9 | sourceforge.net/projects/samtools/files/samtools |

**Table-02.** Software setup installation

| Software | Installation |
| --- | --- |
| JAVA | Follow the installation into the cluster-set up section. |
| WOMtool | Java jar files are accessed over the command-line. |
| CROMWELL |  |
| GATK4 |  |
| SPARK | Follow the installation into the cluster-set up section. |
| HADOOP |  |
| Docker | $ sudo apt-get update  $ sudo apt-get install docker-ce docker-ce-cli containerd.io |
| GATK4 Docker container image | $ docker pull broadinstitute/gatk  $ docker run -it broadinstitute/gatk |
| Bowtie | Export path environment variable to access bowtie executables. |
| Virtual-Box |  |
| Ubuntu OS |  |
| R | Follow installation instructions. |
| RSEM | Compile RSEM using ‘make’ command.  Export path environment variable to access RSEM executables.  Install bioconductor package -  $ R  > source( http://bioconductor.org/biocLite.R) |
| Samtools | $ ./configure  $ make |

1.2 **SNPs**

Mutated genomic bases to call SNPs and their detection status by GATK4 are tabulated in Table-03.

**Table-03.** List of SNPs and their detection status

| Line | Column | Position | Ref | Alt | status |
| --- | --- | --- | --- | --- | --- |
| 39 | 02 | 2592 | A | T | Detected |
| 39 | 09 | 2599 | T | A | Detected |
| 47 | 32 | 3182 | C | G | Detected |
| 88 | 38 | 6058 | T | C | Detected |
| 112 | 18 | 7718 | C | A | Detected |
| 129 | 70 | 8960 | C | A | FP |
| 183 | 21 | 12691 | A | T | Detected |
| 219 | 46 | 15236 | C | G | Detected |
| 240 | 10 | 16670 | A | G | Detected |
| 264 | 20 | 18366 | A | T | FP |

1.3 **INDELs**

Mutated genomic bases to call INDELs and their detection status by GATK4 are tabulated in Table-04.

**Table-04.** List of INDELs and their detection status

| Line | Start Col no | End Col no | Start base position | End base position | Detection status |
| --- | --- | --- | --- | --- | --- |
| 14 | 27 | 40 | 867 | 880 | Not detected |
| 33 | 38 | 68 | 2208 | 2238 | Not detected |
| 58 | 8 | 21 | 3928 | 3941 | Detected |
| 82 | 55 | 60 | 5655 | 5660 | Detected |
| 137 | 31 | 33 | 9481 | 9483 | Detected |
| 191 | 6 | 19 | 13236 | 13249 | Not detected |
| 241 | 35 | 52 | 16905 | 16922 | Not detected |
| 262 | 14 | 34 | 18354 | 18374 | Not detected |
| 266 | 11 | 14 | 18491 | 18494 | Detected |
| 269 | 45 | 45 | 18875 | 18875 | Not detected |

1.4 **TRANSLOCATIONs**

Translocated genomic bases are tabulated in table-05.

**Table-05.** List of TRANSLOCATIONs

|  | **Line** | **Start** | **End** |
| --- | --- | --- | --- |
| 1. | 6 | 1 | 20 |
|  | 33 | 41 | 60 |
| 2. | 45 | 1 | 17 |
|  | 58 | 58 | 274 |
| 3. | 127 | 1 | 18 |
|  | 272 | 60 | 77 |
| 4. | 100 | 1 | 9 |
|  | 113 | 71 | 79 |
| 5. | 117 | 1 | 11 |
|  | 119 | 71 | 81 |

1.5 **INVERSIONs and reverse complement**

Called inversions and their reverse complements in the mutated genome are tabulated in Table-06.

**Table-06.** List of INVERSIONSs

|  | Line | Start | End |
| --- | --- | --- | --- |
| 1. | 120 | 1 | 22 |
| 2. | 129 | 35 | 71 |
| 3. | 233 | 55 | 63 |
| 4. | 41 | 14 | 40 |
| 5. | 55 | 37 | 41 |

Simulated paired-end ebola RNASEQ reads containing above mentioned mutation are deposited at the referenced public repository.(1)

**Table-07.** List of simulated ebola datasets containing non-structural variants

| Mutation type | Mutated fasta file | Left-end reads | Right-end reads |
| --- | --- | --- | --- |
| Non-structural | ebolaMUTANT.fasta | ebolaMUTANT_1.fq | ebolaMUTANT_2.fq |

1.6 **SVs**

There are two datasets containing structural variants - dataset-1 and dataset-2.

Contained structural variants in dataset-1 are tabulated below.

**Table-08.** Dataset-1 and variants detection status

|  | | Dataset - 1 | | | |
| --- | --- | --- | --- | --- | --- |
|  | Mutation type | Start Line | End Line | Total base pair count | Detection  status |
|  | Deletion | 30 | 39 | 700 | Detected |
| Novel sequence insertion | Insertion | 30 | 39 | 700 | Not detected |
|  | Inversion | 30 | 39 | 700 | Not detected |
| Tandem  duplication | Duplication | 30 | 39 | 700 | Not detected |
| Translocated from | Translocation | 30 | 39 | 700 | Detected |
| Translocated to |  | 80 | 89 |  |  |
| CPX1 – INSDUPDEL | Insertion | 30 | 34 | 350 | Detected |
|  | Duplication | 35 | 39 | 350 | Not detected |
|  | Deletion | 45 | 49 | 350 | Detected |
| CPX2 - DELDUPDEL | Deletion | 30 | 34 | 350 | Detected |
|  | Duplication | 35 | 39 | 350 | Not detected |
|  | Deletion | 45 | 49 | 350 | Detected |

Mutated ebola genomes and simulated paired-end RNASEQ reads contained in dataset-1 are deposited to the public repository.(1)

**Table-09.** Dataset-1

| Mutation type | Mutated fasta file | Left-end reads | Right-end reads |
| --- | --- | --- | --- |
| Deletion | ebolaDEL.fasta | ebolaDEL_1.fq | ebolaDEL_2.fq |
| Insertion | ebolaINS.fasta | ebolaINS_1.fq | ebolaINS_2.fq |
| Inversion | ebolaINV.fasta | ebolaINV_1.fq | ebolaINV_2.fq |
| Duplication | ebolaDUP.fasta | ebolaDUP_1.fq | ebolaDUP_2.fq |
| Translocation | ebolaTRAN.fasta | ebolaTRAN_1.fq | ebolaTRAN_2.fq |
| CPX1 - INSDUPDEL | ebolaCPX1.fasta | ebolaCPX1_1.fq | ebolaCPX1_2.fq |
| CPX2 - INSDUPDEL | ebolaCPX2.fasta | ebolaCPX2_1.fq | ebolaCPX2_2.fq |

Contained structural variants in dataset-2 are tabulated below.

**Table-10.** Dataset-2 and detection status

|  | | Dataset -2 | | | |
| --- | --- | --- | --- | --- | --- |
|  | Mutation type | Start Line | End Line | Total base pair count | Detection status |
|  | Deletion1 | 30 | 39 | 700 | Detected |
|  | Deletion2 | 200 | 209 | 700 | Detected |
| Novel  Sequence  insertion | Insertion1 | 30 | 39 | 700 | Not  detected |
|  | Insertion2 | 200 | 209 | 700 | Not  detected |
|  | Inversion1 | 30 | 39 | 700 | Not  detected |
|  | Inversion2 | 200 | 209 | 700 | Not  detected |
| Tandem  duplication | Duplication1 | 30 | 39 | 700 | Not  detected |
|  | Duplication2 | 200 | 209 | 700 | Not  detected |
| Translocated from | Trans-location1 | 30 | 39 | 700 | Detected |
| Translocated to |  | 80 | 89 |  |  |
| Translocated from | Trans-location2 | 200 | 209 | 700 | Detected |
| Translocated to |  | 250 | 259 |  |  |
| CPX1 – INSDUPDEL | Insertion | 30 | 39 | 700 | Not  detected |
|  | Duplication | 40 | 49 | 700 | Not  detected |
|  | Deletion | 50 | 59 | 700 | Detected |
| CPX2 - DELDUPDEL | Deletion | 30 | 39 | 700 | Detected |
|  | Duplication | 40 | 49 | 700 | Not  detected |
|  | Deletion | 50 | 59 | 700 | Detected |

Mutated ebola genomes and simulated paired-end RNASEQ reads contained in dataset-2 are deposited to the public repository.(1)

**Table-11.** Dataset-2

| Mutation type | Mutated fasta file | Left-end reads | Right-end reads |
| --- | --- | --- | --- |
| Deletion | ebolaDEL.fasta | ebolaDEL_1.fq | ebolaDEL_2.fq |
| Insertion | ebolaINS.fasta | ebolaINS_1.fq | ebolaINS_2.fq |
| Inversion | ebolaINV.fasta | ebolaINV_1.fq | ebolaINV_2.fq |
| Duplication | ebolaDUP.fasta | ebolaDUP_1.fq | ebolaDUP_2.fq |
| Translocation | ebolaTRAN.fasta | ebolaTRAN_1.fq | ebolaTRAN_2.fq |
| CPX1 - INSDUPDEL | ebolaCPX1.fasta | ebolaCPX1_1.fq | ebolaCPX1_2.fq |
| CPX2 - INSDUPDEL | ebolaCPX2.fasta | ebolaCPX2_1.fq | ebolaCPX2_2.fq |

1.7 **Setting multi-node SPARK cluster**

The architecture of the cluster has two layers:

- master layer
- slave layer

The entire setup included into the study is done over virtual machines - VirtualBox. Client interacts with the master node at master layer and distributed processing is performed at compute nodes at slave layer. Nodes are connected with each other using computer networking following TCP/IP protocol. All required software needs to be installed over each node of the cluster. Established multi-node SPARK and Hadoop clusters can be used as commodity hardware (to run all SPARK based Computational software).

The Cluster can be set-up using either of the operating systems:

- Red-hat	or
- Debian

All of the hereinafter stated implementation steps are performed over Ubuntu - 16.04 LTS.

### **Make an area for cluster setup**

Make an area over the system, specifying username and password. All of the installation is done into specified areas.

Following is the Linux command-line utility to add user, set password and to obtain root access.

#### *Add user*

- To add/create a new user, follow the command 'useradd' or 'adduser' with <username>.
- The <username> is a user login name that is used by the user to login into the system.

$ useradd <username>	(1)

#### *Set login password*

- To set a login password, the ‘passwd’ command with <username> asks the user to set or change for the login password.

$ passwd <username>	(2)

#### *Obtain root access*

- (Additionally) One may set or change the root password obtaining root access.
- ‘sudo passwd root’ tells the system to change the root password.

$ sudo passwd root	(3)

**Set fully qualified domain name**

- To set a fully qualified domain name (FQDN), mention the IP address of the node, fully qualified domain name and alias for FQDN as tab separated fields onto the /etc/hosts file.

$ sudo su	(4)

$ cat /etc/hosts	(5)

**Examples for setting FQDN for master and slave nodes**

<master_node_ip_address>	master.node.com	master	(6)

<slave_node_ip_address>	slave.node.com	slave	(7)

**Set ssh server and ssh client**

$ ssh_keygen -t rsa -P “”	(8)

- Above command will generate the rsa key. That key is contained into /etc/.ssh/id_rsa file.
- Make its copy into /etc/.ssh/id_rsa.pub as well as into /etc/.ssh/authorized_keys.
- Send this key to all nodes using the scp command-line utility.

$ scp /etc/.ssh/authorized_keys <slave_node_ip_address>:/etc/.ssh/authorized_keys	(9)

- <slave_node_ip_address> is the slave node IP address.
- Do note that all of these changes have to be carried-out as root users*.*

#### *Changes to parameters for root permit login*

- The parameter for root permit login - PermitRootLogin - is contained into /etc/ssh/sshd_config file. *The default setting for the parameter is <prohibit password>*. Change it to yes.
- Run the following command-line utility to save the changes:

$ service sshd restart	(10)

#### *Setting JAVA environment for the run-time*

- Get root access.
- Make a sub-directory /java/ into /usr/lib/ directory.
- Copy your extracted jdk folder into /usr/lib/java.
- Set full permission to java directory.

$ sudo su	(11)

$ chmod 777 /usr/lib/java	(12)

- Access .bashrc file & add the following block at the end of that file:

$ export JAVA_HOME=/usr/lib/java/<jdk-setup-name>	(13)

$ export PATH=”$PATH:$JAVA_HOME/bin”	(14)

- Make these environment variables and path variables permanent over the system after updating the .bashrc file.

$ source .bashrc	(15)

- Run the java installation command over command-line.

$sudo update-alternatives --install "/usr/bin/java" "java" "/usr/lib/java/<jdk-setup-name>/bin/java" 1	(16)

$sudo update-alternatives --install "/usr/bin/javac" "javac" "/usr/lib/java/<jdk-setup-name>/bin/javac" 1	(17)

$sudo update-alternatives --install "/usr/bin/javaws" "javaws" "/usr/lib/java/<jdk-setup-name>/bin/javaws" 1	(18)

1.8 **SPARK installation**
- Get root access.
- Make a sub-directory */spark/* into */usr/lib/* directory.
- Copy extracted spark setup into the */usr/lib/spark* directory.
- Make the following changes to the .bashrc file.

$ export SPARK_HOME=/usr/lib/spark/<spark-setup-name>	(19)

$ export PATH=”$PATH:$SPARK_HOME/bin”	(20)

- Make these environment variables and path variables permanent over the system after updating the .bashrc file.

$ source .bashrc	(21)

- As spark runs on top of the java environment, make the following changes to configure spark with java.
  - Navigate to the *SPARK_HOME/conf* directory.
  - Copy spark-env-templet.sh to spark-env.sh.

$ cd $SPARK_HOME/conf	(22)

$ cp slave-env-templet.sh slave-env.sh	(23)

- To the spark-env.sh file, mention java installation path by adding the following block to it:

$ export JAVA_HOME=/usr/lib/java/<jdk-setup-name>	(24)

**Introduction of IP addresses to slave.templet**

- Copy slave.templet file that is present in the *$SPARK_HOME/conf* directory.

$ cd $SPARK_HOME/conf	(25)

$ cp slave.templet slave	(26)

- Mention IP addresses of the nodes to the copied file.

<master_node_ip_address>	(27)

<slave_node_ip_address>	(28)

- Check for java and spark installation.

$ echo $JAVA_HOME	(29)

$ echo $SPARK_HOME	(30)

- Both of the aforementioned commands will return the absolute path of installed set-up for JAVA and SPARK.
- Start SPARK services over your node.

$ sudo su	(31)

$ ./$SPARK_HOME/sbin/start-all.sh	(32)

- Access it over web-browser using localhost:7077.

Running SPARK-based GATK4 tools in cluster mode will require the Hadoop native library. The installation of Hadoop is mentioned hereinafter.

1.9 **HADOOP installation**
- Create a new directory */usr/lib/hadoop* onto both machines.
- Extract, copy and paste hadoop set-up folders and files inside */user/lib/hadoop* directory.
- Access the .bashrc file and add the following block at the end of the file:

export HADOOP_HOME=/usr/lib/hadoop/hadoop-2.6.5	(33)

export HADOOP_INSTALL=$HADOOP_HOME	(34)

export HADOOP_MAPRED_HOME=$HADOOP_HOME	(35)

export HADOOP_COMMON_HOME=$HADOOP_HOME	(36)

export HADOOP_HDFS_HOME=$HADOOP_HOME	(37)

export YARN_HOME=$HADOOP_HOME	(38)

export HADOOP_COMMON_LIB_NATIVE_DIR=$HADOOP_HOME/lib/native	(39)

export PATH=$PATH:$HADOOP_HOME/sbin:$HADOOP_HOME/bin	(40)

export HADOOP_CONF_DIR=$HADOOP_HOME/etc/hadoop	(41)

- Update the .bashrc file to save all of the made changes by the following command:

$source ~/.bashrc	(42)

- If one wishes to establish or integrate full-fledged Hadoop cluster, one may go for editing or changing the configurations or parameters of following configuration files in order to change port number, number of CPU cores and HDFS daemons files:
  - core-site.xml
  - hdfs-site.xml
  - mapred-site.xml
  - yarn-site.xml
  1.10 **Writing WDL scripts**

All of the WDL scripts as well as their corresponding input files are deposited to the public repository.(1)

**Table-12.** List of all workflows and WDL scripts

| Workflow | WDL script | Input file |
| --- | --- | --- |
| SNPs and INDELs  detection using non-SPARK based GATK4 tools. | variantcall.wdl | variantcall_inputs.json |
| SNPs and INDELs  detection using SPARK based GATK4 tools. | variantcallSPARK.wdl | variantcallSPARK_inputs.json |
| SNPs and INDELs detection using non-SPARK based GATK4 tools and Docker as run-time environment. | variantcallDOCKER.wdl | variantcallDOCKER_inputs.json |
| SNPs and INDELs  detection using SPARK based GATK4 tools and Docker as run-time  environment. | variantcallSPARKDOCKER.wdl | variantcallSPARKDOCKER_inputs.json |
| Structural-variants  detection using SPARK based GATK4 tools. | structuralvariantSPARK.wdl | structuralvariantSPARK_inputs.json |
| Structural-variants  detection using SPARK based GATK4 tools and Docker as run-time  environment. | structuralvariantSPARKDOCKER.wdl | structuralvariantSPARKDOCKER_inputs.json |

1.11 **WDL script for “SNPs and INDELs detection using SPARK based GATK4 tools and Docker as run-time environment”**

The description or anatomy of WDL script for “SNPs and INDELs detection using SPARK based GATK4 tools and Docker as run-time environment” is as follows:

### **workflow variantcall → Workflow block workflow level**

#### *Input variables*

- File refIndex → Samtools generated reference genome index file
- File refDict → Reference genome PICARD dictionary file
- File referenceGenome → Reference genome file
- String gatk_docker → Docker container image
- String gatk_path → Path of gatk jar file (placed within the docker container)
- String name → String value that serves as the base name for specified file (especially o/p file)

### **call Alignment → Tasks call and enumeration between workflow level and task level**

#### *Input variables*

- ReferenceGenome=referenceGenome
- sampleName=name
- index=name

### **call AddOrReplaceReadGroups**

#### *Input variables*

- inputSAM=Alignment.rawSAM → Linear chaining
- docker=gatk_docker
- gatk_path=gatk_path
- sampleName=name

### **call SortSamSpark**

#### *Input variables*

- inputBAM=AddOrReplaceReadGroups.rawBAM → Linear chaining
- sampleName=name
- docker=gatk_docker
- gatk_path=gatk_path

### **call ReferenceSeqIndex**

#### *Input variables*

- ReferenceGenome=referenceGenome
- sampleName=name

### **call ReferenceSeqDictionary**

#### *Input variables*

- docker=gatk_docker
- gatk_path=gatk_path
- ReferenceGenome=referenceGenome
- sampleName=name

### **call MarkDuplicatesSpark**

#### *Input variables*

- inputBAM=SortSamSpark.rawBAM → Linear chaining
- docker=gatk_docker
- gatk_path=gatk_path
- sampleName=name

### **call SplitNCigarReads**

#### *Input variables*

- inputBAM=MarkDuplicatesSpark.rawBAM → Linear chaining
- RefIndex=refIndex
- RefDict=refDict
- docker=gatk_docker
- gatk_path=gatk_path
- ReferenceGenome=referenceGenome
- sampleName=name

### **call HaplotypeCallerSpark**

#### *Input variables*

- inputBAM=SplitNCigarReads.rawBAM → Linear chaining
- RefIndex=refIndex
- RefDict=refDict
- docker=gatk_docker
- gatk_path=gatk_path
- sampleName=name

### **call VariantFilteration**

#### *Input variables*

- mutantVCF=HaplotypeCallerSpark.rawVCF → Linear chaining
- RefIndex=refIndex
- RefDict=refDict
- docker=gatk_docker
- gatk_path=gatk_path
- ReferenceGenome=referenceGenome
- sampleName=name

### **call SelectSNPs**

#### *Input variables*

- mutantVCF=VariantFilteration.rawVCF → Linear chaining
- docker=gatk_docker
- gatk_path=gatk_path
- ReferenceGenome=referenceGenome
- RefIndex=refIndex
- RefDict=refDict
- sampleName=name

### **call SelectINDELs**

#### *Input variables*

- mutantVCF=VariantFilteration.rawVCF → Linear chaining
- docker=gatk_docker
- gatk_path=gatk_path
- ReferenceGenome=referenceGenome
- RefIndex=refIndex
- RefDict=refDict
- sampleName=name

### **task Alignment → Alignment task block**

#### *Input variables*

- File leftFastq → left-end fastq file
- File rightFastq → right-end fastq file
- File ReferenceGenome → Reference genome file
- String sampleName → String value serves as base name for specified files
- String index → String value for base name for index

#### *command → Command block*

export PATH=$PATH:/absolute/path/of/bowtie-executables/	(43)

bowtie2-build ${ReferenceGenome} ${index}	(44)

bowtie2 -q -x ${index} -1 ${leftFastq} -2 ${rightFastq} -S ${sampleName}.sam	(45)

#### *output → Output block*

- File rawSAM = "${sampleName}.sam" → Output file

### **task ReferenceSeqIndex → Task block for samtools index**

#### *Input variables*

- File ReferenceGenome → Reference genome file.
- String sampleName → String value for sample name

#### *command → Command block*

samtools faidx ${ReferenceGenome} > ${sampleName}.fasta.fai	(46)

#### *output → Output block*

- File refIndex = "${sampleName}.fasta.fai"	→ Output file

### **task ReferenceSeqDictionary → Task block for CreateSequenceDictionary**

#### *Input variables*

- String docker → String value for docker container image
- String gatk_path → Path of gatk jar file (placed within the docker container)
- File ReferenceGenome → Reference genome file
- String sampleName → String value for sample name

#### *command → Command block*

java -jar ${gatk_path} CreateSequenceDictionary -R ${ReferenceGenome} -O ${sampleName}.fasta.dict	(47)

#### *runtime → Runtime block*

- docker:docker

#### *output → Output block*

- File refDict = "${sampleName}.fasta.dict" → Output file

### **task AddOrReplaceReadGroups → Task block for AddOrReplaceReadGroups**

#### *Input variables*

- String docker → String value for docker container image
- String gatk_path → Path of gatk jar file (placed within the docker container)
- File inputSAM → Input SAM file generated by Alignment
- String sampleName → String value for sample name

#### *command → Command block*

java -jar ${gatk_path} AddOrReplaceReadGroups -I ${inputSAM} -O ${sampleName}.bam -RGID 1 -RGLB 445_LIB -RGPL illumina -RGSM RNA -RGPU illumina	(48)

#### *runtime → Runtime block*

- docker:docker

#### *output → Output block*

- File rawBAM = "${sampleName}.bam" → Output file

### **task SortSamSpark → Task block for SortSam**

#### *Input variables*

- String docker → String value for docker container image
- String gatk_path → Path of gatk jar file (placed within the docker container)
- File inputBAM → Input BAM file generated by AddOrReplaceReadGroups
- String sampleName → String value for sample name

#### *command → Command block*

java -jar ${gatk_path} SortSamSpark -I ${inputBAM} -O ${sampleName}.bam	(49)

#### *runtime → Runtime block*

- docker:docker

#### *output → Output block*

- File rawBAM = "${sampleName}.bam" → Output file

### **task MarkDuplicatesSpark → Task block for MarkDuplicates**

#### *Input variables*

- String docker → String value for docker container image
- String gatk_path → Path of gatk jar file (placed within the docker container)
- File inputBAM → Input BAM file generated by SortSam
- String sampleName → String value for sample name

#### *command → Command block*

java -jar ${gatk_path} MarkDuplicatesSpark -I ${inputBAM} -O ${sampleName}.markdup.bam -M output.metrics	(50)

#### *runtime → Runtime block*

- docker:docker

#### *output → Output block*

- File rawBAM = "${sampleName}.markdup.bam" → Output file name

### **task SplitNCigarReads → Task block for SplitNCigarReads**

#### *Input variables*

- String docker → Docker container image
- String gatk_path → Path of gatk jar file (placed within the docker container)
- File inputBAM → Input BAM file generated by MarkDuplicatesSpark
- File ReferenceGenome → Reference genome file
- File RefIndex → Reference genome index file generated by samtools index.
- File RefDict → Dictionary file of reference genome
- String sampleName → String value for sample name

#### *command → Command block*

java -jar ${gatk_path} SplitNCigarReads -R ${ReferenceGenome} -I ${inputBAM} -O ${sampleName}.split.bam	(51)

#### *runtime → Runtime block*

- docker:docker

#### *output → Output block*

- File rawBAM = "${sampleName}.split.bam"

### **task HaplotypeCallerSpark → Task block for HaplotypeCaller**

#### *Input variables*

- String docker → Docker container image
- String gatk_path → Path of gatk jar file (placed within the docker container)
- File inputBAM → Input BAM file generated by SplitNCigarReads
- File Reference2bitGenome → 2bit format reference genome file
- File RefIndex → Reference genome samtools index file
- File RefDict → Dictionary file of reference genome
- String sampleName → String value for sample name

#### *command → Command block*

samtools index ${inputBAM} > ${sampleName}.split.bam.bai	(52)

java -jar ${gatk_path} HaplotypeCallerSpark -R ${Reference2bitGenome} -I ${inputBAM} -O ${sampleName}.mutant.vcf	(53)

#### *runtime → Runtime block*

- docker:docker

#### *output → Output block*

- File rawVCF = "${sampleName}.mutant.vcf" → Output file

### **task VariantFilteration → Task block for VariantFiltration**

#### *Input variables*

- String docker → Docker container image
- String gatk_path → Path of gatk jar file (placed within the docker container)
- File mutantVCF → VCF file generated by HaplotypeCallerSpark
- File ReferenceGenome → Reference genome file
- File RefIndex → Reference genome samtools index file
- File RefDict → Dictionary file of reference genome
- String sampleName → String value for sample name

#### *command → Command block*

java -jar ${gatk_path} IndexFeatureFile -F ${mutantVCF}	(54)

java -jar ${gatk_path} VariantFiltration -R ${ReferenceGenome} -V ${mutantVCF} -window 35 -cluster 3 -O ${sampleName}.mutantfilter.vcf	(55)

#### *runtime → Runtime block*

- docker:docker

#### *output → Output block*

- File rawVCF = "${sampleName}.mutantfilter.vcf" → Output file name

### **task SelectSNPs → Task block for SelectVariants**

#### *Input variables*

- String docker → Docker container image
- String gatk_path → Path of gatk jar file (placed within the docker container)
- File mutantVCF → VCF file generated by VariantFiltration
- File ReferenceGenome → Reference genome file
- File RefIndex → Reference genome samtools Index file
- File RefDict → Dictionary file of reference gnome
- String sampleName → String value for sample name

#### *command → Command block*

java -jar ${gatk_path} SelectVariants -R ${ReferenceGenome} -V ${mutantVCF} -O ${sampleName}.mutantsnp.vcf -select-type-to-include SNP	(56)

#### *runtime → Runtime block*

- docker:docker

#### *output → Output block*

- File rawVCF = "${sampleName}.mutantsnp.vcf" → Output file

### **task SelectINDELs → Task block for SelectVariants**

#### *Input variables*

- String docker → Docker container image
- String gatk_path → Path of gatk jar file (placed within the docker container)
- File mutantVCF → VCF file generated by VariantFiltration
- File ReferenceGenome → Reference genome file
- File RefIndex → Reference genome samtools Index file
- File RefDict → Dictionary file of reference genome
- String sampleName → String value for sample name

#### *command → Command block*

java -jar ${gatk_path} SelectVariants -R ${ReferenceGenome} -V ${mutantVCF} -O ${sampleName}.mutantindel.vcf -select-type-to-include INDEL	(57)

#### *runtime → Runtime block*

- docker:docker

#### *output → Output block*

- File rawVCF = "${sampleName}.mutantindel.vcf" → Output file

The input file for the WDL script is in JSON format file containing key-value pairs. Their description or anatomy is as follows:

- "variantcall.name": "ebola" → Sample name
- "variantcall.refDict": "/path of directory/ebola.dict" → Reference genome dictionary file
- "variantcall.refIndex": "/path of directory/ebola.fasta.fai" → Reference genome samtools index file
- "variantcall.Alignment.rightFastq": "/path of directory/ebolaMUTANT_2.fq" → Right-end fastq reads
- "variantcall.referenceGenome": "/path of directory/ebola.fasta" → Reference genome file
- "variantcall.gatk_docker":"broadinstitute/gatk:4.1.0.0"	→	GATK4 docker image
- "variantcall.gatk_path": "/gatk/gatk.jar" → Path of gatk jar file placed within its container
- "variantcall.Alignment.leftFastq":	"/path	of	directory/ebolaMUTANT_1.fq" → Left-end fastq reads
- "variantcall.HaplotypeCallerSpark.Reference2bitGenome": "/path of directory/ebola.fasta" → Reference genome in 2bit format
1.12 **WDL script for SV detection using SPARK-based GATK4 tools and Docker as run-time environment**

The description or anatomy of WDL script for SV detection using SPARK-based GATK4 tools and Docker as run-time environment is as follows:

**workflow variantcall → Workflow block workflow level**

#### *Input variables*

- String gatk_path → Path of gatk jar file (placed within docker container)
- String gatk_docker → Docker container image
- File refIndex → Reference genome Index file
- File refDict → Dictionary file of reference genome
- File referenceGenome → Reference genome file
- String name → String value for sample name

### **call Alignment → Task call and enumeration between workflow level and task level input variables**

#### *Input variables*

- ReferenceGenome = referenceGenome
- sampleName = name
- index = name

### **call AddOrReplaceReadGroups**

#### *Input variables*

- inputSAM=Alignment.rawSAM → Linear chaining
- sampleName=name
- docker=gatk_docker
- gatk_path=gatk_path

### **call SortSam**

#### *Input variables*

- inputBAM = AddOrReplaceReadGroups.rawBAM → Linear chaining
- docker = gatk_docker
- gatk_path = gatk_path
- sampleName = name

### **call ReferenceSeqIndex**

#### *Input variables*

- ReferenceGenome = referenceGenome
- sampleName = name

### **call ReferenceSeqDictionary**

#### *Input variables*

- docker = gatk_docker
- gatk_path = gatk_path
- ReferenceGenome = referenceGenome
- sampleName = name

### **call BwaMemIndexImageCreator**

#### *Input variables*

- docker = gatk_docker
- gatk_path = gatk_path
- ReferenceGenome = referenceGenome
- sampleName = name

### **call FindBadGenomicKmers**

#### *Input variables*

- docker = gatk_docker
- gatk_path = gatk_path
- ReferenceGenome = referenceGenome
- RefDict = refDict
- RefIndex = refIndex
- sampleName = name

### **call FindBreakpointEvidence**

#### *Input variables*

- docker = gatk_docker
- gatk_path = gatk_path
- ReferenceGenome = referenceGenome
- inputBAM = SortSam.rawBAM → Linear chaining
- image = BwaMemIndexImageCreator.indexImage → Linear chaining
- KmersToIgnore = FindBadGenomicKmers.kmersToIgnore	→	Linear chaining
- RefDict = refDict
- sampleName = name

### **call SvDiscovery**

#### *Input variables*

- docker = gatk_docker
- gatk_path = gatk_path
- ReferenceGenome = referenceGenome
- inputSAM = FindBreakpointEvidence.rawSAM → Linear chaining
- RefDict = refDict
- RefIndex = refIndex
- sampleName = name

### **task Alignment → Task block for alignment**

#### *Input variables*

- File leftFastq → Left-end fastq reads
- File rightFastq → Right-end fastq reads
- File ReferenceGenome → Reference genome file
- String sampleName → String value for sample name
- String index → String value for indexes

#### *command → Command block*

export PATH=$PATH:/absolute/path/of/bowtie_executables	(58)

bowtie2-build ${ReferenceGenome} ${index}	(59)

bowtie2 -q -x ${index} -1 ${leftFastq} -2 ${rightFastq} -S ${sampleName}.sam	(60)

#### *output → Output block*

- File rawSAM = "${sampleName}.sam"	→ Output file name

### **task ReferenceSeqIndex → Task block for reference index**

#### *Input variables*

- File ReferenceGenome → Reference genome file name
- String sampleName → String value for sample name

#### *command → Command block*

samtools faidx ${ReferenceGenome} > ${sampleName}.fasta.fai	(61)

#### *output → Output block*

- File refIndex = "${sampleName}.fasta.fai" → Output file

### **task ReferenceSeqDictionary → Task block for CreateSequenceDictionary**

#### *Input variables*

- String docker → Docker container image
- String gatk_path → Path of gatk jar file (placed within docker container)
- File ReferenceGenome → Reference genome file
- String sampleName → String value for sample name

#### *command → Command block*

java -jar ${gatk_path} CreateSequenceDictionary -R ${ReferenceGenome} -O ${sampleName}.fasta.dict	(62)

#### *runtime → Runtime block*

- docker:docker

#### *output → Output block*

- File refDict = "${sampleName}.fasta.dict" → Output file name

### **task AddOrReplaceReadGroups → Task block for AddOrReplaceReadGroups**

#### *Input variables*

- String docker → Docker container image
- String gatk_path → Path of gatk jar file (placed within docker container)
- File inputSAM → Input SAM file generated by Alignment
- String sampleName → String value for sample name

#### *command → Command block*

java -jar ${gatk_path} AddOrReplaceReadGroups -I ${inputSAM} -O ${sampleName}.bam -RGID 1 -RGLB 445_LIB -RGPL illumina -RGSM RNA -RGPU illumina	(63)

#### *runtime → Runtime block*

- docker:docker

#### *output → Output block*

- File rawBAM = "${sampleName}.bam"	→ Output file name

### **task SortSam → Task block for SortSam**

#### *Input variables*

- String docker → Docker container image
- String gatk_path → Path of gatk jar file (placed within docker container)
- File inputBAM → Input BAM file generated by AddOrReplaceReadGroups
- String sampleName → String value for sample name

#### *command → Command block*

java -jar ${gatk_path} SortSam -I ${inputBAM} -O ${sampleName}.bam -CREATE_INDEX true -VALIDATION_STRINGENCY LENIENT -SO coordinate	(64)

#### *runtime → Runtime block*

- docker:docker

#### *output → Output block*

- File rawBAM = "${sampleName}.bam" → Output file

### **task BwaMemIndexImageCreator → Task block for BwaMemIndexImageCreator**

#### *Input variables*

- String docker → Docker container image
- String gatk_path → Path of gatk jar file (placed within docker container)
- File ReferenceGenome → Reference genome file
- String sampleName → String value for sample name

#### *command → Command block*

java -jar ${gatk_path} BwaMemIndexImageCreator -I ${ReferenceGenome} -O ${sampleName}.img	(65)

#### *runtime → Runtime block*

- docker:docker

#### *output → Output block*

- File indexImage = "${sampleName}.img" → Output file

### **task FindBadGenomicKmers → Task block for FindBadGenomicKmers**

#### *Input variable*

- String docker → Docker container image
- String gatk_path → Path of gatk jar file (placed within docker container)
- File ReferenceGenome → Reference genome file
- String sampleName → String value for sample name
- String kmerstoignore → String value kmerstoignore
- File RefDict → Dictionary file generated by ReferenceSeqDictionary
- File RefIndex → Index file generated by ReferenceSeqIndex

#### *command → Command block*

java -jar ${gatk_path} FindBadGenomicKmersSpark -R ${ReferenceGenome} -O ${kmerstoignore}.txt	(66)

#### *runtime → Runtime block*

- docker:docker

#### *output → Output block*

- File kmersToIgnore = "${kmerstoignore}.txt" → Output file

### **task FindBreakpointEvidence → Task block for FindBreakpointEvidence**

#### *Input variables*

- String docker → Docker container image
- String gatk_path → Path of gatk jar file (placed within docker container)
- File inputBAM → Input BAM file generated by SortSam
- File ReferenceGenome → Reference genome file
- File KmersToIgnore → KmersToIgnore generated by FindBadGenomicKmers
- File image → Image file generated by BwaMemIndexImageCreator
- File RefDict → Dictionary file generated by ReferenceSeqDictionary
- String sampleName → String value for sample name

#### *command → Command block*

java -jar ${gatk_path} FindBreakpointEvidenceSpark -I ${inputBAM} --aligner-index-image ${image} --kmers-to-ignore ${KmersToIgnore} -O ${sampleName}.assemblies.sam	(67)

#### *runtime → Runtime block*

- docker:docker

#### *output → Output block*

- File rawSAM = "${sampleName}.assemblies.sam" → Output file

### **task SvDiscovery → Task block for SvDiscovery**

#### *Input variables*

- String docker → Docker container image
- String gatk_path → Path of gatk jar file (placed within docker container)
- File inputSAM → Input SAM file generated by FindBreakpointEvidence
- String sampleName → String value for sample name
- File RefDict → Dictionary file generated by ReferenceSeqDictionary
- File RefIndex → Index file generated by ReferenceSeqIndex
- File ReferenceGenome → Reference genome file

#### *command → Command block*

java -jar ${gatk_path} CreateSequenceDictionary -R ${ReferenceGenome} -O ${sampleName}.dict	(68)

java -jar ${gatk_path} SvDiscoverFromLocalAssemblyContigAlignmentsSpark -I ${inputSAM} -R ${ReferenceGenome} -O ${sampleName}	(69)

#### *runtime → Runtime block*

- docker:docker

#### *output → Output block and output files*

- File rawDICT="${sampleName}.dict"
- File rawOUTPUT1="${sampleName}_RNA_NonComplex.vcf"
- File rawOUTPUT2="${sampleName}_RNA_Complex.vcf"
- File	rawOUTPUT3="${sampleName}_RNA_cpx_reinterpreted_simple_1_seg.vcf"
- File	rawOUTPUT4="${sampleName}_RNA_cpx_reinterpreted_simple_multi_seg.vcf"
- File rawOUTPUT5="${sampleName}_RNA_merged_simple.vcf"

The description or anatomy of the input json file for the above mentioned structural variant discovery WDL script is as follows:

- "variantcall.name": "ebola" → Sample name
- "variantcall.refDict": "/path to directory/ebola.fasta.dict" → Reference genome dictionary file
- "variantcall.refIndex": "/path to directory/ebola.fasta.fai" → Reference genome samtools index file
- "variantcall.gatk_docker": "broadinstitute/gatk:4.1.0.0" → GATK4 docker image
- "variantcall.FindBadGenomicKmers.kmerstoignore": "kmers_to_ignore" → String value for kmerstoignore
- "variantcall.Alignment.rightFastq": "/path to directory/ebolaDEL_2.fq" → Right-end fastq reads
- "variantcall.referenceGenome": "/path to directory/ebola.fasta" → Reference genome fasta file
- "variantcall.gatk_path": "/gatk/gatk.jar" → Path of gatk jar file placed within it`s container
- "variantcall.Alignment.leftFastq": "/path to directory/ebolaDEL_1.fq" → Left-end fastq reads

**References**

1.	KUMAR A. Implementation of Genomic Variant Calling Using GATK4, SPARK, WDL, CROMWELL and DOCKER Over Simulated Ebola NGS Dataset. 2020 May 23;
